# Aligning single-cell developmental and reprogramming trajectories identifies molecular determinants of reprogramming outcome

**DOI:** 10.1101/122531

**Authors:** Davide Cacchiarelli, Xiaojie Qiu, Sanjay Srivatsan, Michael Ziller, Eliah Overbey, Jonna Grimsby, Prapti Pokharel, Ken Livak, Shuqiang Li, Alex Meissner, Tarjei Mikkelsen, John Rinn, Cole Trapnell

**Affiliations:** Armenise-Harvard laboratory of Integrative Genomics, Telethon Institute of Genetics and Medicine, Pozzuoli, Italy; Department of Stem Cell and Regenerative Biology, Harvard University, Cambridge, Massachusetts, USA; The Broad Institute of MIT and Harvard, Cambridge, Massachusetts, USA; Harvard Stem Cell Institute, Harvard University, Cambridge, Massachusetts, USA; Molecular & Cellular Biology Program, University of Washington, Seattle, WA, USA; Department of Genome Sciences, University of Washington, Seattle, WA, USA.; Max-Planck Institute of Psychiatry, Munich, Germany; Present address: Dana-Farber Cancer Institute, Boston, MA; Fluidigm Corporation, South San Francisco, CA, USA

## Abstract

Cellular reprogramming through manipulation of defined factors holds great promise for large-scale production of cell types needed for use in therapy, as well as for expanding our understanding of the general principles of gene regulation. MYOD-mediated myogenic reprogramming, which converts many cell types into contractile myotubes, remains one of the best characterized model system for direct conversion by defined factors. However, why MYOD can efficiently convert some cell types into myotubes but not others remains poorly understood. Here, we analyze MYOD-mediated reprogramming of human fibroblasts at pseudotemporal resolution using single-cell RNA-Seq. Successfully reprogrammed cells navigate a trajectory with two branches that correspond to two barriers to reprogramming, with cells that select incorrect branches terminating at aberrant or incomplete reprogramming outcomes. Differential analysis of the major branch points alongside alignment of the successful reprogramming path to a primary myoblast trajectory revealed Insulin and BMP signaling as crucial molecular determinants of an individual cell’s reprogramming outcome, that when appropriately modulated, increased efficiency more than five-fold. Our single-cell analysis reveals that MYOD is sufficient to reprogram cells only when the extracellular milieu is favorable, supporting MYOD with upstream signaling pathways that drive normal myogenesis in development.

## Introduction

During development, cells undergo drastic shifts in gene expression and epigenetic configuration as they pass from progenitor or stem cell states to their differentiated states in the adult organism. Nevertheless, developmental decisions can be “unmade” by ectopic expression of a small number of regulatory genes, as first shown by Davis et al, who converted murine fibroblasts to myotubes by overexpressing MyoD, a key myogenic transcription factor (Davis et al., 1987). Numerous other direct reprogramming factors have since been discovered, most famously including four factors that reprogram many different cell types into pluripotent stem cells (iPSCs)(Takahashi and Yamanaka, 2006). Following the discovery of MyoD-mediated transdifferentiation in fibroblasts, Weintraub et. al. explored MyoD’s ability to convert a diverse set of starting cell types to myotubes (Weintraub et al., 1989). Curiously, ectopic expression of MyoD induced the expression of muscle specific structural proteins without altering cell specific functions. These studies, concomitant with contemporary heterokaryon experiments (Blau et al., 1983), indicated that neither intracellular transcription factors or ectopically transduced MyoD were sufficient for complete reprogramming to the myogenic cell fate. Despite the following decades of research on cellular conversions, including extensive studies of myogenic reprogramming, the molecular determinants that impede or mediate reprogramming remain poorly characterized.

Reprogramming can generate dramatically heterogeneous cell populations, with cells reaching different drastically different molecular outcomes. Single-cell genomics assays provide a means of identifying the different outcomes and could reveal mechanisms that drive a cell to a particular one. For example, Truetlein et al. demonstrated through a single-cell transcriptome analysis that overexpression of Ascl1, Brn2 and Myt1l generates both induced neurons and an alternative, myocyte-like cell fate (Treutlein et al., 2016). However, synthesizing measurements from these diverse outcomes into a useful picture of the molecular mechanisms determine cell fate remains extremely challenging. Recently, we developed an algorithm that can automatically reconstruct the sequence of expression changes executed by a cell undergoing differentiation or reprogramming from single-cell RNA-Seq data. Our algorithm, called Monocle, introduced the notion of “pseudotime”, which measures each cell’s progress through a biological process without the need for a priori knowledge of genes that define progression (Trapnell et al., 2014). Moreover, Monocle can pinpoint branches that lead a cell to alternative outcomes, which can reveal the genes that direct a cell to its ultimate fate. Monocle’s unsupervised learning algorithm enables the discovery of key steps, roadblocks, and intermediate cellular states on the path to differentiation.

Here, we apply our pseudotemporal analysis approach to analyze human fibroblasts undergoing MYOD-mediated myogenic reprogramming. Through an unsupervised single-cell RNA-Seq analysis, we reveal that the trajectory includes branch points corresponding to key “checkpoints” in the process. Cells that travel down the correct branch progress, while those that travel down the alternatives fail to fully convert to myotubes. We developed a novel approach for aligning two pseudotime trajectories, which showed that the myogenic conversion trajectory does contain a path that is strikingly similar to normal myoblast differentiation, but that many cells diverge from this path toward unproductive reprogramming outcomes. Detailed inspection of genes differentially regulated at these branch points revealed that insulin and BMP signaling are major molecular determinants of myogenic conversion. Modulation of either of these pathways alone substantially improved conversion efficiency, and this effect was potentiated by modulating them together. Pseudotemporal analysis is thus a powerful technique for isolating the sequence of productive state transitions leading to effective cell type conversion.

## Results

### Pseudotemporal analysis of MYOD overexpression reveals multiple reprogramming outcomes

To analyze myogenic reprogramming at single-cell resolution, we first derived a doxycycline-inducible MYOD fibroblast line (hFib-MyoD), enabling us to synchronize the onset of MYOD expression. The hFib-MyoD line also harbors a constitutively expressed human telomerase gene to alleviate passage-dependent stress, which we previously showed drastically improves uniformity and synchrony of induced pluripotency across cells (Cacchiarelli et al., 2015). 72 hours of *MYOD* transgene expression was sufficient to produce conversion of fibroblasts to myosin heavy chain-positive (MYH+) cells. However, as reported by previous studies (Salvatori et al., 1995), myogenic reprogramming of human fibroblasts is a very inefficient process. Only a small proportion of cells are successfully reprogrammed, and the result are small, mononucleated myocyte-like cells (Figure 1A,B) rather than long, multinucleated myotubes as generated by human skeletal myoblasts (HSMM). The number of reprogrammed cells did not increase with longer time courses (not shown).

To investigate the molecular basis driving HSMM and hFIB-MYOD through the process of myogenic identity, we sampled single cells every 24 hours post myogenic induction and performed deep, single-cell, full-length RNA sequencing. Although some markers of myotube formation, such as enolase 3 (a.k.a skeletal muscle enolase; *ENO3*) were induced by MYOD, others including dystrophin (DMD) and the myogenic transcription factor myogenin (*MYOG*) showed expression patterns inconsistent with HSMM (Figure 1C). Few hFib-MYOD cells expressed MYOG even at 72 hours, which is required for the upregulation of many genes needed for terminal differentiation and contraction, including *DMD*(Bentzinger et al., 2012)*. CDK1* and other genes associated with active proliferation, which were rapidly downregulated in HSMM within 24 hours following switch to differentiation medium, were more gradually lost, with some cells still expressing them even at 72 hours. Delayed cell cycle exit may explain why hFib-MYOD wells contained greater than 3-fold more cells than HSMM wells despite being seeded at the same initial density (Figure 1B). Comparing average expression levels in cells collected at each time point revealed that few of the 653 genes that were significantly differentially expressed as a function of time in myoblasts were regulated to the same extent in hFib-MYOD (Figure 1D).

**Figure 1.**
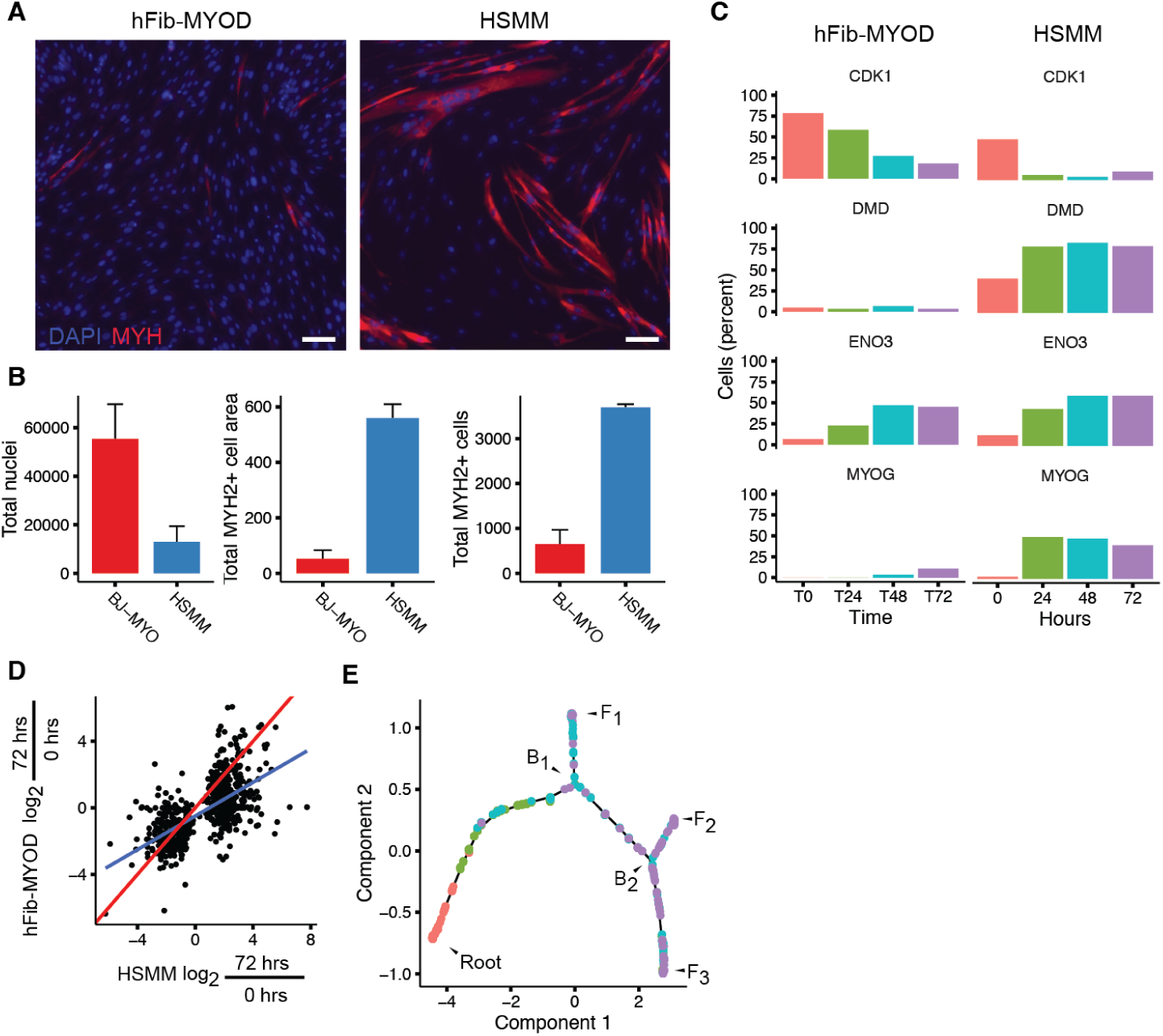
hFib-MYOD is a human fibroblast line that inefficiently converts to myotube-like cells upon doxycycline (dox) induction. Equal numbers of both cells were plated at identical density (25000/cm2). **A**) Immunostaining of hFib-MYOD and HSMM with muscle-specific anti-myosin-heavy chain (a-MYH2) antibodies 72 hours post induction of MYOD-mediated reprogramming or myoblast differentiation via serum switch, respectively. Scale bar is 100um. **B**) Counts of MYH2+ cells, size of MYH2+ cells in pixels, and total nuclei as measured by automated image processing scripts (see Methods). **C**) Fraction of cells in which mRNAs for selected muscle markers were detected via full-length single-cell RNA-Seq. **D**) Fold changes in average expression level of genes significantly differentially expressed (FDR < 5%) between 0 and 72 hours in differentiating myoblasts compared to their corresponding changes in hFib-MYOD. The blue line indicates a linear regression, while the red line illustrates perfect concordance. **E**) The single-cell trajectory reconstructed by Monocle 2 for hFib-MYOD cells undergoing myogenic reprogramming. Cells start at the root and progress to one of three alternative reprogramming outcomes, denoted F_1_, F_2_, and F_3_. To reach these fates, cells must pass through branch point B_1_. Cells that do not proceed to F_1_ must then chose between F_2_ or F_3_ at branch point B_2_.

### Single-cell trajectory branch points correspond to reprogramming barriers

We next sought to identify the subset of hFib-MYOD cells that reached a muscle-like expression program and define the stepwise changes in gene expression that lead to conversion failure. Since both differentiation and reprogramming are characterized by high levels of asynchrony, obscuring the sequence of expression changes in such processes, we applied a recently improved version of Monocle(Qiu et al., 2017a) to hFib-MYOD. Monocle 2 reconstructed a trajectory capturing the progression of single cells through myogenic reprogramming, (Figure 1F) which contained three termini (denoted “F_1_”, “F_2_”, and “F_3_”) corresponding to three distinct reprogramming outcomes. To reach these outcomes, cells passed through at least one of two branch points (denoted “B_1_” and “B_2_”). The trajectory’s root was populated exclusively by cells collected at the beginning of the experiment, while the three termini of the tree were populated by cells collected following the switch to differentiation media. Reanalysis of our previously collected HSMM data using a similar approach with Monocle 2 uncovered a branch point in the trajectory, leading to one of two differentiation outcomes (Qiu et al., 2017a).

**Figure 2.**
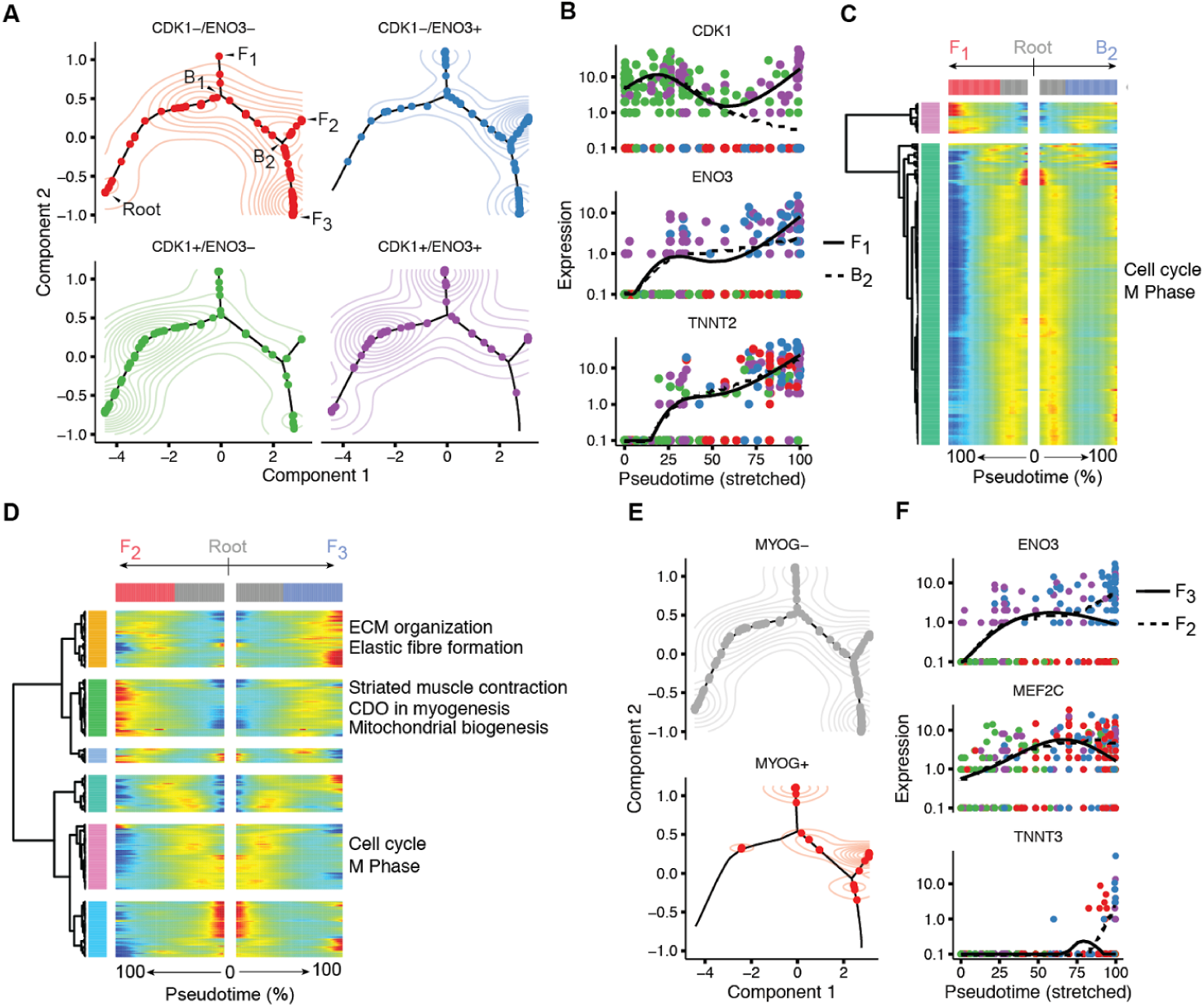
Single cell trajectory analysis with Monocle 2 identifies three MYOD-mediated myogenic reprogramming outcomes. A) Cells along the trajectory divided into four groupings based on expression of CDK1 and ENO3. Contour plots indicate two-dimensional gaussian kernel density estimates. B) Pseudotime kinetics of CDK1, ENO3, and TNNT2 from the root of the trajectory to outcome F1 (solid line) and the cells up to branch point B_2_. C) All genes expressed in a branch B_1_-dependent manner. Each row indicates the standardized (row-centered) kinetic curves of a gene. The center of the heatmap shows the kinetic curve value at the root of the trajectory. Proceeding from the center to the left of the heatmap follows the kinetic curve from the root along the trajectory to outcome F_1_. Proceeding to the right yields the curve from the root to B_2_. D) Cells colored by detected of MYOG mRNA. **E**) Kinetic curves for ENO3, MEF2C, and TNNT3 from the root through B_2_ to outcomes F_2_ (dashed line) and F_3_. F) Branched kinetic heatmap for all genes dependent on B_2_, along with selected enriched gene ontology terms in indicated clusters.

We next assessed the expression of genes regulated in myoblast differentiation in cells at the three different outcomes of the hFib-MYOD trajectory. Surprisingly, cells at outcome F_1_ showed expression of genes upregulated early in myoblast differentiation, such as *ENO3*, but maintained strong expression of genes needed for active proliferation such as *CDK1*. (Figure 2A) In contrast, during differentiation of HSMMs, key genes that mark proliferating cells (*CCNB2* and *CDK1*) are almost never co-expressed with early myogenic markers (*ENO3* and *TNNT1*) (Supplemental Figure 1). Although hFib-MYOD cells at F_1_ co-expressed numerous markers of both proliferation and the early myoblast differentiation program, they expressed few markers of mature, contractile myotubes. We next examined pseudotemporal kinetics of *CDK1, ENO3*, and *TNNT2* as cells traveled from the root through branch point B_1_ and then either to outcome F_1_ or towards a second branch point B_2_. While *ENO3* and *TNNT2* were upregulated to a similar extent on both paths, *CDK1* levels dropped markedly in cells traveling to the branch B_2_, but remained high in cells traveling to F_1_ (Figure 2B). A global differential analysis comparing the two paths away from branch point B_1_ revealed 173 genes with branch-dependent expression (FDR < 1%), most of which were associated with the cell cycle. (Figure 2C) One path leading away from branch point B_1_ reached a second branch point B_2_, which in turn gave rise to outcomes F_2_ and F_3_. Branch-dependent expression analysis at B_2_ showed significant changes in 277 genes (FDR < 10%), including a coherent cluster that included *TNNI1, TNNT3, MYL4*, and other notable proteins involved in myotube contraction. (Figure 2D) Cells at F_3_ expressed *MYOG*, lacked markers of active proliferation, and showed upregulation of numerous genes needed for muscle contraction (Figure 2E-F, Supplemental Figure 2). In contrast, cells at F_2_ lacked expression of *MYOG*, cell cycle genes, and most of those needed for muscle contraction. Based on our differential analysis, we termed the three hFib-MYOD reprogramming F_1_, F_2_, and F_3_ outcomes as “cell cycle exit failure”, “partial reprogramming”, and “reprogramming failure”, respectively (Figure 3A).

**Figure 3.**
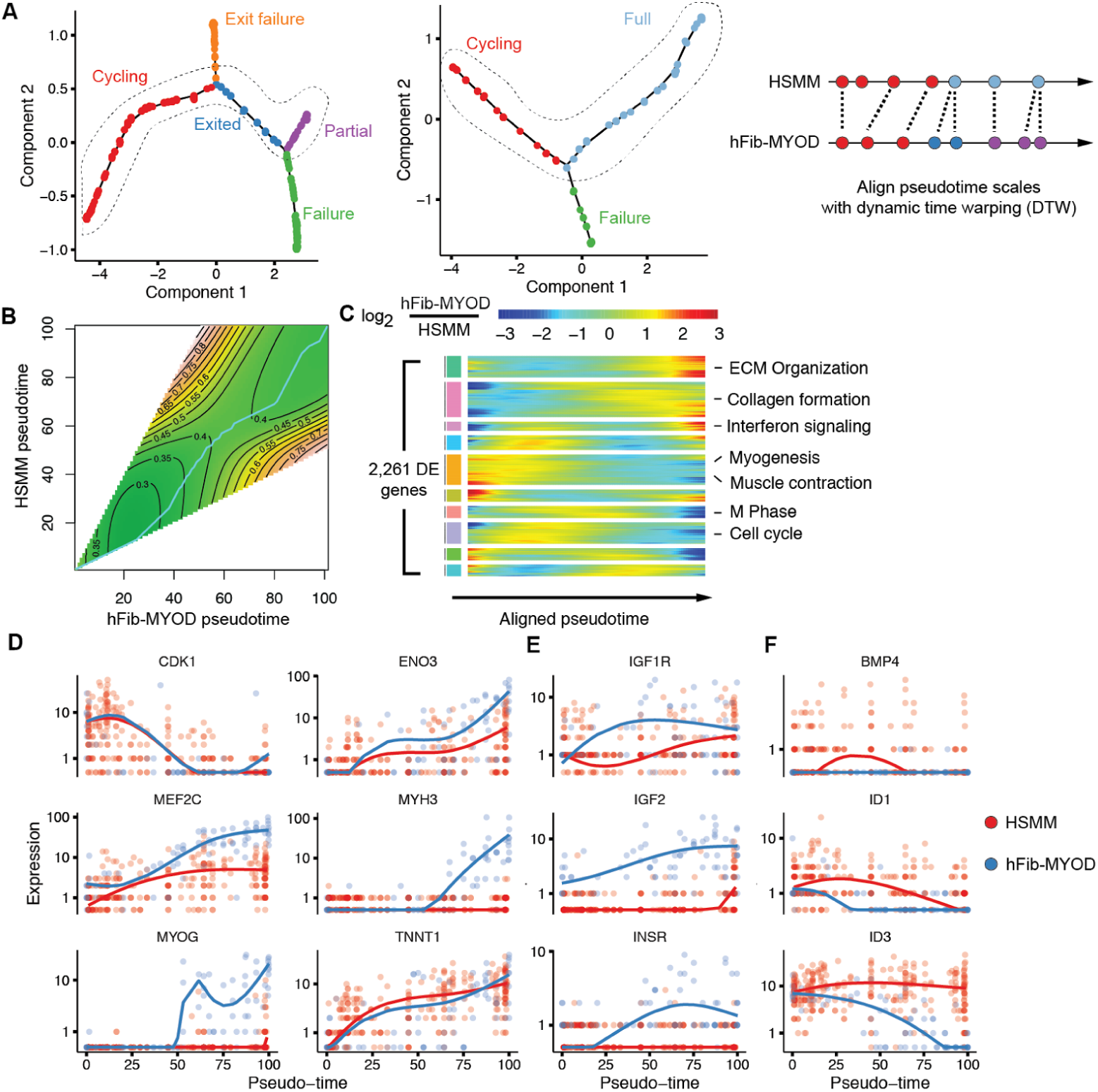
Alignment of myogenic reprogramming and myoblast differentiation trajectories identifies insulin and BMP signaling as aberrantly regulated in hFib-MYOD. **A**) The core trajectories of hFib-MYOD (left) and HSMM (center) were aligned using Dynamic Time Warping (DTW), a dynamic-programming time series alignment technique, which matches pseudotime coordinates on the hFib-MYOD trajectory to the most best matching points in the HSMM trajectory as determined by similarity in global expression. **B**) The DTW alignment (cyan curve) follows a dissimilarity-minimizing path through the “landscape” of possible ways to pair up points on the two pseudotime curves. The contours indicate levels of equal similarity between global expression profiles of typical cells at indicated HSMM and hFib-MYOD pseudotimes. **C**) Clustered heatmap of genes showing significant differences in aligned kinetic curves. Each shows the log-transformed fold change of the curve for hFib-MYOD over that of HSMM at each point in the aligned trajectory. Genes are clustered by Ward’s method and tested for enrichment of genes in REACTOME pathways, with selected significantly enriched pathways shown. **D**) Aligned kinetic curves for several key indicators of myogenic progress. Each row is a gene reported as significantly different between HSMM and hFib-MYOD (FDR < 10%; likelihood ratio test; Methods) after controlling for common pseudotime-dependent differences. Values shown are the instantaneous log_2_-transformed fold changes between spline-smoothed expression curves computed separately for each cell type. These values were further transformed into per-gene Z-scores prior to clustering. **E**) Aligned kinetic curves for IGF1R, IGF2, and INSR, and **F**) BMP4, ID1, and ID3, all of which show significant differential pseudotime-dependent expression.

### Aligning single-cell trajectories identifies key molecular determinants of myogenic reprogramming

Having identified the core trajectory from the fibroblast state to a partially reprogrammed state, we next sought to quantify the differences between this program of gene expression changes and the one associated with normal myoblast differentiation. Because the two single-cell trajectories were learned from each dataset independently, their pseudotime scales are not directly comparable; there is no universal “unit” of pseudotime. We thus devised an alignment algorithm based on Dynamic Time Warping (DTW) (Vintsyuk, 1968) which matches highly similar points on the two trajectories and creates a mapping between the HSMM and hFib-MYOD pseudotime scales. (Figure 3A-C). The alignment revealed extensive quantitative differences in expression kinetics between the two trajectories, highlighting several coherent clusters of genes missing in hFib-MYOD or aberrantly regulated relative to HSMM (Figure 3D). Most of the cell-cycle markers, such as CDK1, displayed nearly identical kinetics in both hFib-MYOD and HSMM, indicating that on the core hFib-MYOD trajectory, the cell cycle exit proceeds swiftly following media switch. MEF2C, a crucial co-factor for MYOD and MYOG in myoblasts, was upregulated with similar timing, but to a lesser extent in hFib-MYOD. MYOG, in contrast, was upregulated far later in hFib-MYOD than HSMM, to a lesser extent, and in a smaller proportion of cells. (Figure 3E)

In myoblasts, MEF2C and MYOG both auto-regulate their own expression (Edmondson et al., 1992; Wang et al., 2001), after initially being upregulated by MYOD. MYOD activates its targets by forming a heterodimer with E47, which together recruit p300 to robustly transactivate the late myogenic expression program. Activity of p300 at MYOD-bound promoters is driven through Akt-mediated insulin signaling (Serra et al., 2007). Formation of the MYOD/E47 heterodimer is inhibited by ID family proteins (Neuhold and Wold, 1993), which sequester E proteins away from chromatin. In proliferating myoblasts ID family proteins are maintained at high levels by an incompletely understood mechanism that is likely downstream of BMP signaling (Lewis and Prywes, 2013), which acts to allow sufficient myoblast expansion from satellite cells during regeneration (Sartori et al., 2013). During differentiation, ID expression drops, enabling E47 to form heterodimers with MYOD at promoters of key muscle genes. Expression levels of insulin receptor (INSR) and the insulin-like growth factor receptors (IGF1R and IGF2R) were significantly lower in hFib-MYOD. (Figure 3F) IGF2, which drives myoblast differentiation in an autocrine loop (Florini et al., 1991), was strongly upregulated in HSMMs but not hFib-MYOD. Furthermore, ID1 and ID3 were at far higher levels in hFib-MYOD, and were maintained throughout differentiation. BMP4 was expressed in hFib-MYOD, while HSMMs did not express BMP family mRNAs at appreciable levels. (Figure 3G) We thus hypothesized that high levels of BMP and insufficient insulin signaling were locking hFib-MYOD cells in a negative feedback loop, preventing their efficient activation of the myotube expression program.

### Cell-extrinsic factors are required for MYOD-mediated myogenic reprogramming

To test whether modulating Insulin or BMP signaling could enhance myogenic conversion efficiency, we supplemented differentiation medium with recombinant Insulin protein, a chemical inhibitor of the BMP receptor, or both. After 120 hours of conversion in the presence of Insulin, hFib-MYOD showed marked increases in the number of myotubes. (Figure 4A) Myoblasts cultured with Insulin did not form significantly more myotubes, but myotube size, the number of nuclei residing in myotubes and nuclei per myotube all increased at least two-fold, indicating far more efficient fusion. Inhibition of BMP signaling also significantly increased the number, size, and nuclei count of hFib-MYOD-derived myotubes. (Figure 4B) The addition of both Insulin and the BMP inhibitor increased myotube number, size, and nucleation more than adding either alone, suggesting that these pathways act independently to modulate hFib-MYOD conversion efficiency. Importantly, neither Insulin nor the BMP inhibitor altered the total number of nuclei in the experiment, indicating that variation in myotube efficiency was not due to variation in cell counts. Together, these results suggest that MYOD is sufficient to convert human foreskin fibroblasts to myotubes only in the presence of appropriate upstream signaling from core myogenic pathways.

**Figure 4.**
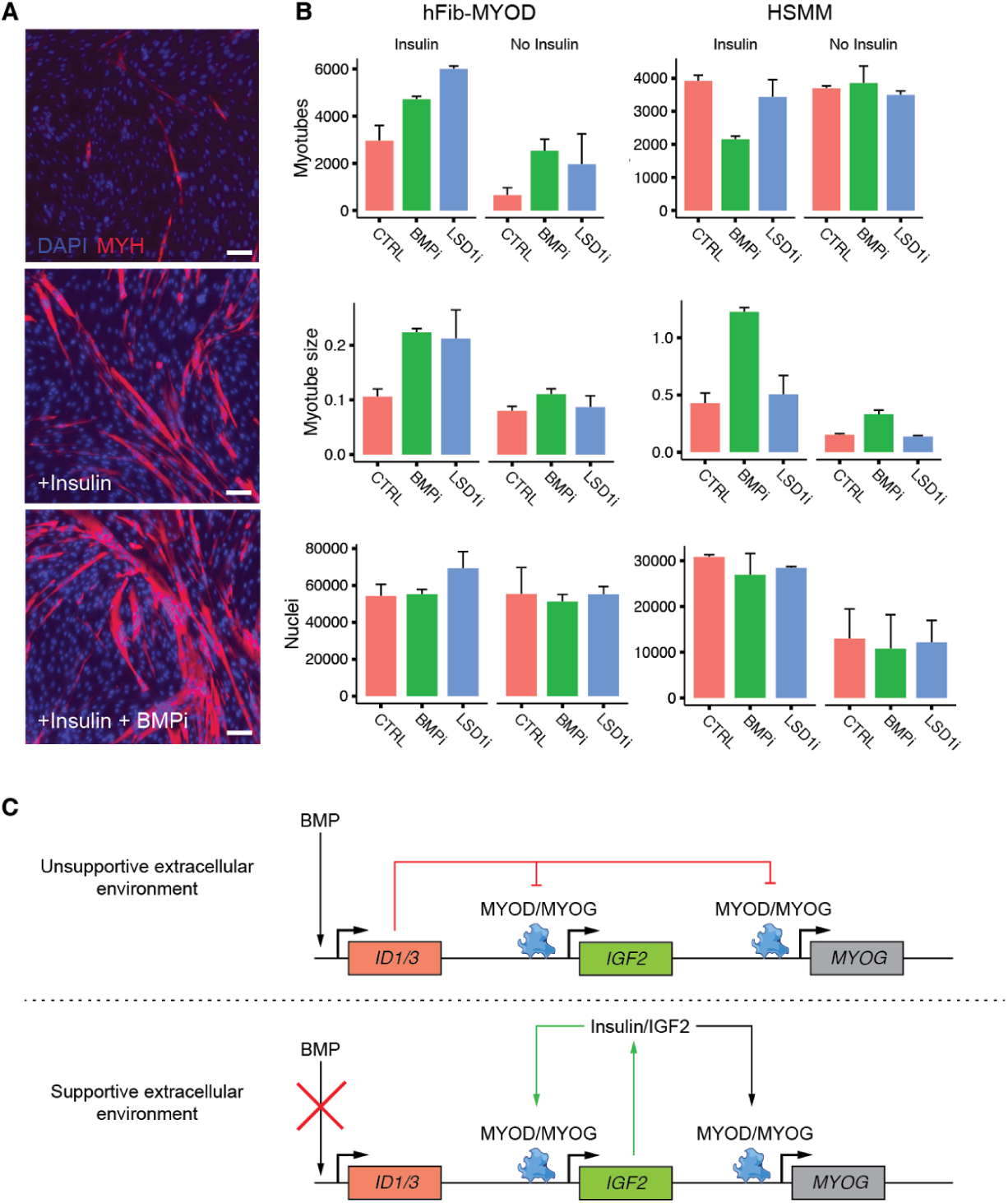
Adding Insulin, inhibiting BMP signaling, and inhibiting LSD1 activity increase myogenic conversion efficiency. **A**) Immunostaining of hFib-MYOD with muscle-specific anti-myosin-heavy chain (a-MYH2) antibodies 72 hours post induction of MYOD-mediated reprogramming. **B**) Counts of MYH2+ cells, size of MYH2+ cells in pixels, and total nuclei as measured by automated image processing scripts (see Methods) **C**) A model summarizing how the extracellular signaling environment conditions cells for reprogrammability by MYOD. An unsupportive environment that exposes cells to BMP maintains cellular ID family protein levels, which in turn suppress the myogenic program by interfering with formation of active MYOD-containing transcription complexes. A supportive environment lacking BMP and containing Insulin engages an IGF2 mediated autocrine feedback loop that amplifies expression of *MYOG* and potentially its co-factors, leading to efficient induction of the myogenic expression program.

MYOD has been characterized as a “pioneer” transcription factor because of its sufficiency in remodeling chromatin at inactive regulatory sites in the genome (Zaret and Carroll, 2011). Recently, a number of studies have shown that genetically or chemically modulating the activity of chromatin remodeling enzymes alters the efficiency, timing, and heterogeneity of reprogramming to the pluripotent state. For example, we showed that inhibiting activity of the histone demethylase LSD1, which removes mono- and di-methyl groups from lysine 4 in the tail of histone H3 (H3K4m1/2), improves fibroblast-to-iPS conversion efficiency by tenfold (Cacchiarelli et al., 2015). Similarly, incubating hFib-MYOD cells with an LSD1 inhibitor dramatically increased their conversion to myotubes in the presence of insulin (Figure 4B). In contrast, the inhibitor had only a modest effect in the absence of Insulin, and no effect on HSMM differentiation.

## Discussion

The landmark discovery by Takahashi and Yamanaka that developmental decisions can be reversed with the ectopic expression of four factors, ultimately converting fibroblasts into pluripotent cells, raised the prospect of large scale manufacture of arbitrary cell types and tissues. (Takahashi and Yamanaka, 2006). During reprogramming, several key developmental decisions are “unmade” in a reverse-stepwise fashion (Cacchiarelli et al.,2015; Takahashi et al., 2014). However, for most cell types, we lack effective protocols for differentiating them from pluripotent cells or direct “reprogramming cocktails” that would generate them from another cell type. The cocktails that do exist generally only work on one or a handful of initial cell types, often with poor and variable efficiency (Vierbuchen and Wernig, 2011), (Xu et al., 2015). Developing better methods for dissecting the molecular basis for different reprogramming cocktails is therefore critical for realizing the promise of efficient manufacture of therapeutically relevant cells.

MYOD is perhaps the best characterized ectopic reprogramming factor, and has been extensively studied as a model “master regulator” of cell fate. Central to its identity is its ability to singlehandedly convert various cell types into muscle. However, as noted some cell types, such as HeLa cells(Weintraub et al., 1989), remain refractory to myogenic conversion for reasons that until recently remained poorly understood. Work by Forcales et. al. showed that HeLa cells can be converted to myotubes with MYOD only when BAF60C, a subunit of the BAF ATP-dependent chromatin remodeling complex (present in mesodermal lineages) is also expressed along with appropriate signaling from p38α (Forcales et al., 2012). That is, MYOD’s ability to reprogram to muscle is dependent upon the presence of factors for needed for ATP-dependent chromatin remodeling.

Our study shows that the extracellular signaling environment must also be supportive for MYOD-mediated reprogramming. Using pseudotemporal single-cell transcriptome analysis, we show that MYOD alone is not sufficient to engage a central autocrine positive feedback loop driven by Insulin signaling (mainly through secreted IGF2 in myoblasts). Moreover, human foreskin fibroblasts secrete BMP4, which may engage a negative autocrine loop and impedes myogenic conversion. Supplementing the cellular milieu with Insulin and a BMP inhibitor rescued MYOD’s ability to convert them to myotubes. Dependence on a supportive extracellular signaling environment might explain variable reprogramming efficiency during not only myogenic reprogramming but many other reprogramming settings. Furthermore, while the importance of engaging regulatory feedback loops for reprogramming, differentiation, and maintenance of the pluripotent and other cellular states is well appreciated, feedback loops are most frequently discussed in the context of transcriptional circuits that govern these processes. Our work underscores the importance of establishing (or interrupting) autocrine signaling feedback loops in order to reach a desired cellular state.

Detailed analysis of the single-cell trajectories for reprogrammed fibroblasts and myoblasts pinpointed these molecular barriers to myogenic conversion. These trajectories contained branches, revealed by a recently improved version of our pseudotime ordering algorithm Monocle, that corresponded to key regulatory barriers to myoblast differentiation and reprogramming. Trajectory branches have been reported to correspond to cell-fate decisions, but also arise in response to genetic perturbations. Identifying genes that differ between the two paths leading away from a branch point can reveal the regulatory mechanisms by which cells decide which path to take. (Qiu et al., 2017a, 2017b). The MYOD reprogramming trajectory contained a core path that resembled normal myoblast differentiation, but quantitatively aligning it to the myoblast differentiation program revealed that key myogenic regulators downstream of insulin and BMP signaling were incompletely activated even in cells on this path.

Beyond providing insights into barriers to myogenic reprogramming, our study shows that pseudotemporal single-cell trajectory analysis with Monocle is a powerful approach for dissecting biological processes with multiple outcomes. Trajectories enable two complementary types of comparative analyses. The first is branch differential analysis, which compares the expression changes that lead to one fate against the changes that lead to another. The second is trajectory alignment, in which two distinct biological processes are compared at pseudotime resolution. This can reveal kinetic differences in how each gene is regulated in the two processes, and could be useful for understanding not only how reprogramming compares to normal cell differentiation, but how a dynamic process proceeds differently under different environmental conditions or how a mutation alters it. Branch analysis and trajectory alignment with Monocle are both applicable to these and many other settings, and should yield insights into the molecular mechanisms of reprogramming, cell differentiation, and a wide array of other biological processes.

## Supplemental Figures

**Supplementary Figure 1.**
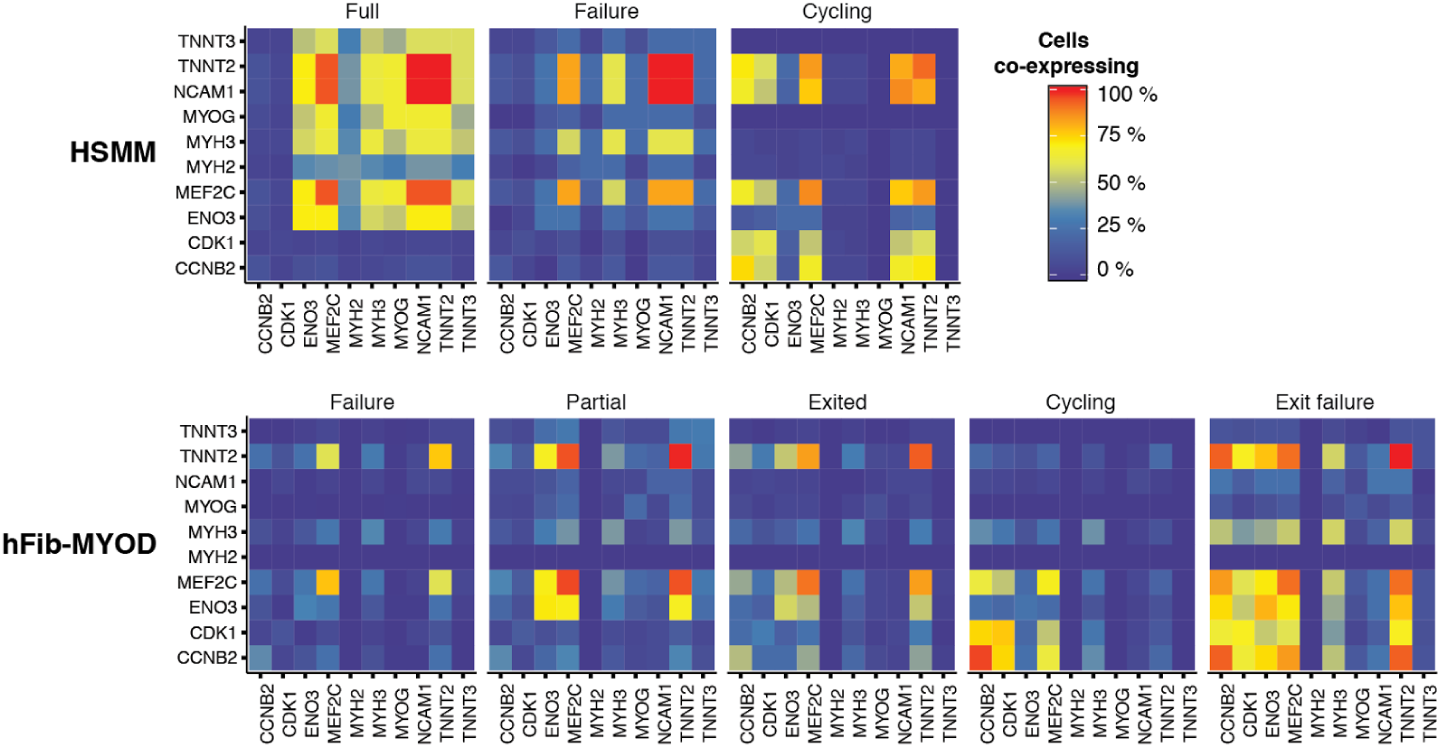
Fraction of cells co-expressing selected markers of proliferation and myoblast differentiation for both HSMM and hFib-MYOD cells.

**Supplementary Figure 2.**
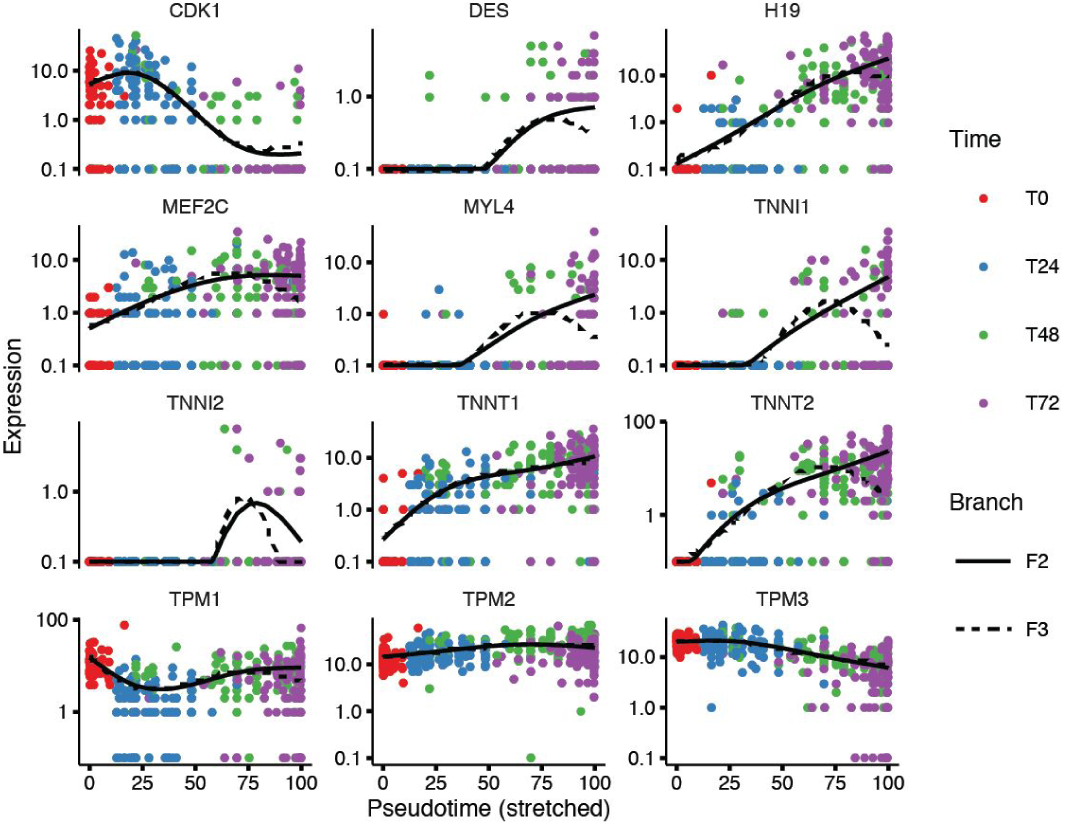
Pseudotemporal expression kinetics for selected markers of proliferation and myoblast differentiation as hFib-MYOD cells transit from the root to outcomes F_2_ or F_3_. Cell’s progressing to F_1_ are excluded for clarity.

## Methods

### Cell culture and derivation

HSMM derivation, expansion and differentiation was as previously described (Trapnell et al., 2014). Foreskin Human Fibroblasts obtained from commercial vendor (Stemgent) were expanded in alpha-mem supplemented with glutamax, 10% FBS and 16ng/ul of FGF-b (ThermoFisher Scientific). The fibroblasts were then infected with a mixture of lentiviruses encoding hTERT (and Puromycin resistance gene), Tetracycline Repressor (and Geneticin resistance gene) and TR-controlled hMYOD (and Blasticidin resistance gene) (ThermoFisher Scientific). All the resistance genes are constitutively expressed and therefore triple selection was performed and maintained using 1ug/ml of Puromycin, 500ug/ml of Geneticin and 2ug/ml of Blasticidin to generate the hFib-MYOD line.

To perform reprogramming and differentiation experiments HSMM and hFib-MYOD cells were plated in 24-well formats at a density of 50.000 cells per well. Gelatin was sometime used as a coating agent with no significant difference with respect to uncoated dishes. Differentiation and reprogramming was induced using a differentiation media containing alpha-mem supplemented with glutamax and 2% HS (ThermoFisher Scientific), supplemented with 2ug/ml of doxycycline to enact MYOD expression. When indicated, Insulin was used at 8ug/ml (ThermoFisher Scientific), BMP inhibitor LDN-193189 was used at 0.1uM (Stemgent), LSD1 inhibitor RN-1 was used at 1uM (EMD Millipore).

### Imaging-based quantification of myotube formation

At the indicated time points, cells were fixed, stained, imaged and processed as previously described(Trapnell et al., 2014). Briefly, we fixed the cells in 4% PFA, stained for and MHC antibody at a 1:500 dilution (ebioscience) and whole-well imaged using a Celigo Microscope (Nexcelom). The imaging/counting script is available upon request.

### Single-cell transcriptome sequencing

At the indicated time points, cells were harvested by gentle dissociation using TrypLE (ThermoFisher Scientific) and processed for full-length transcriptome sequencing using the Fluidigm C1 Single Cell or bulk mRNA sequencing as previously described (Trapnell et al., 2014). Cells were loaded onto the microfluidic chip at a concentration of ∽250 cells/μl to maximize the number of single cells captured and minimize the occurrence of doublets. In some instances, cells were reloaded if the majority of capture sites were not occupied on the first attempt. As in Trapnell et al, 2014, captured cells were scored by manual on-chip microscopic inspection to determine if they were singletons and free of other debris. Gene expression profiles for each cell were computed using TopHat (v 2.1.1) and Cufflinks (2.2.0) software packages, which were provided with the hg19 genomic reference and GENCODE v17 transcriptome index.

### Single-cell trajectory reconstruction

Single-cell pseudotime trajectories were constructed with Monocle version 2.2.0. Monocle 2 is a dramatically more powerful method than our previous software. Briefly, it uses Reversed Graph Embedding, a machine learning technique that given high dimensional data, constructs a principal graph approximating a manifold in a low-dimensional space. Single cells are projected onto this manifold, which orders them into a trajectory and identifies any branch points corresponding to cell fate decisions. (Qiu et al., 2017a)

Myoblasts were ordered as in Qiu et al. Briefly, expression levels were first converted into relative mRNA counts using Census (Qiu et al., 2017b). Cells with more than 1e6 Census RNAs were filtered as cells that failed conversion. From this set, cells with +/- 2 standard deviations more or less than the mean total Census RNAs were also excluded as either low-complexity libraries or suspected doublets not caught by visual inspection. The HSMM culture contains a mixture of myoblasts and contaminating interstitial fibroblasts, so we informatically classified each cell on the basis of semi-supervised clustering. We then collected a set of ordering genes that defined myoblast differentiation by testing each gene for differential expression between the four time points in the experiment. The top 100 most significant genes were selected for downstream ordering analysis. Expression profiles were reduced to 2 dimensions using the DDRTree algorithm (Mao et al) included with Monocle 2, via the `reduceDimension ()`, with `ncenter=50` **and** `param. gamma=100`.

hFib-MYOD cells were filtered for quality and ordered similarly to the myoblasts, though we made no attempts to split the population, as there is no contamination of other cells types in the culture. hFib-MYOD cells were ordered on the basis of the same genes used to order the myoblasts, excluding those genes not detected in at least 5 cells. As there are more hFib-MYOD cells than myoblasts, parameters to `reduceDimension ()`, were adjusted (`ncenter=100` **and** `param. gamma=100`).

### Trajectory alignment

We use Dynamic Time Warping (DTW) to compute the optimal alignment between the HSMM and hFib-MYO trajectories. We refer readers to (Giorgino and Others, 2009; Rabiner and Juang, 1993)for a detailed discussion, and summarize the algorithm below. Briefly, DTW, aims to compare two longitudinal data series to one another by locally stretching or compressing them and make them resemble each other while preserving their order. The algorithm takes as input a distance matrix *d* that captures the dissimilarity between each pair of elements of a vector *X* and *Y*. Because DTW takes as input the matrix *d*, rather than directly examining *X* and *Y*, they can be vectors of multivariate observations, categorical values, or some mix of these, provided that the user defines a suitable distance function. DTW computes a “warping curve” *ø*(*k*), *k* = 1, …,*T:*

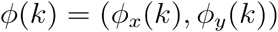

Where the warping function *ø*_*x*_ maps the time indices of *X* to that of *Y* and and *ø*_*y*_ maps *Y* indices to that of *X*. DTW then can compute the alignment cost of a particular warping curve as

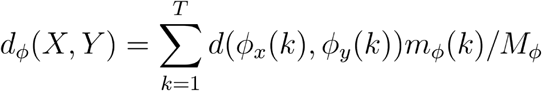

The per-step weight *m*_*ø*_(*k*)and the normalization constant *Mø* aim to make the accumulated distortions along different alignment paths comparable. DTW then finds optimal alignment *ø* (subject to additional constraints to preserve the ordering of the data points) via dynamic programming.

To align the hFib-MYO and HSMM trajectories, we first selected cells along the “productive” paths from the root of each trajectory to outcomes F_2_ and F_1_, respectively. Next we generated smoothed expression curves (using Monocle 2’s genSmoothCurves() function) for the union of the ordering genes used to create each cell type’s trajectory. Genes for which curves could not be generated due to numerical instabilities in either cell type were excluded. These curves were then standardized independently, scaling each to have a mean of zero and a standard deviation of 1. The curves were collected into a pair of matrices, *H* and *M* for the myoblasts and the hFib-MYO cells, respectively, with one row per gene and one column for each of the 100 points on each gene’s smoothed curve. The pairwise dissimilarities between each of the 100 pseudo time points in *M* and the 100 points in *F* were then computed as *d*(*i, j*) = 1 – *cor*(*H_i_, M_j_*). These pairwise distances capture the global similarity between myoblasts and hFib-MYO cells at each pair of points on their trajectories. DTW allows the user to specify the cost of skipping points in one time series or another, analogously to how dynamic programming for sequence alignment imposes different costs for insertions, deletions, and mismatches. Here, we provide DTW with a “step pattern” specified by `rabinerJuangStepPattern (3, “d”)`. To project the two trajectories onto a common pseudotime scale, we use the `warp ()` function of the `dtw` package.

### Differential expression analysis

We tested for differential gene expression as a function of pseudotime (*Ψ*) as previously described (Qiu et al., 2017b; Trapnell et al., 2014): We fit the following model

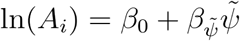

Where A_*i*_ is the mean of a negative-binomial valued random variable of the Census-normalized transcript count for gene *i*, and the tilde above *Ψ* indicates that these predictors are smoothed with natural splines during fitting. This model was compared to the reduced model:

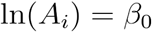
by likelihood ratio test. Genes with an Benjamini-Hochberg adjusted p-value of less than 0.05 were determined to be dynamic across pseudotime.

When testing for differences between HSMM and hFib-MYOD, we used a modified model to identify genes that were regulated in a pseudotime-dependent manner differently between the two cell types:

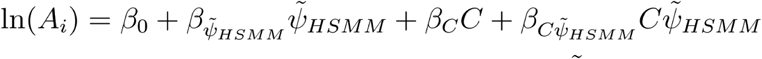

Where C is an indicator variable encoding the cell type, and 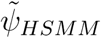 is the pseudotime scale of HSMM, onto which the hFib-MYOD cells were projected by DTW. This full model was compared to

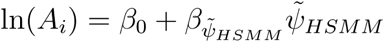

By likelihood ratio test to identify genes with cell-type dependent differences in expression, while controlling for differences in expression due to pseudotime that might be common to the two.

## References

Bentzinger, C.F., Wang, Y.X., and Rudnicki, M.A. (2012). Building muscle: molecular regulation of myogenesis. Cold Spring Harb. Perspect. Biol. 4.

Blau, H.M., Chiu, C.P., and Webster, C. (1983). Cytoplasmic activation of human nuclear genes in stable heterocaryons. Cell 32, 1171–1180.

Cacchiarelli, D., Trapnell, C., Ziller, M.J., Soumillon, M., Cesana, M., Karnik, R., Donaghey, J., Smith, Z.D., Ratanasirintrawoot, S., Zhang, X., et al. (2015). Integrative Analyses of Human Reprogramming Reveal Dynamic Nature of Induced Pluripotency. Cell 162, 412–424.

Davis, R.L., Weintraub, H., and Lassar, A.B. (1987). Expression of a single transfected cDNA converts fibroblasts to myoblasts. Cell 51, 987–1000.

Edmondson, D.G., Cheng, T.C., Cserjesi, P., Chakraborty, T., and Olson, E.N. (1992). Analysis of the myogenin promoter reveals an indirect pathway for positive autoregulation mediated by the muscle-specific enhancer factor MEF-2. Mol. Cell. Biol. 12, 3665–3677.

Florini, J.R., Magri, K.A., Ewton, D.Z., James, P.L., Grindstaff, K., and Rotwein, P.S. (1991). “Spontaneous” differentiation of skeletal myoblasts is dependent upon autocrine secretion of insulin-like growth factor-II. J. Biol. Chem. 266, 15917–15923.

Forcales, S.V., Albini, S., Giordani, L., Malecova, B., Cignolo, L., Chernov, A., Coutinho, P., Saccone, V., Consalvi, S., Williams, R., et al. (2012). Signal-dependent incorporation of MyoD-BAF60c into Brg1-based SWI/SNF chromatin-remodelling complex. EMBO J. 31, 301–316.

Giorgino, T., and Others (2009). Computing and visualizing dynamic time warping alignments in R: the dtw package. J. Stat. Softw. 31, 1–24.

Lewis, T.C., and Prywes, R. (2013). Serum regulation of Id1 expression by a BMP pathway and BMP responsive element. Biochim. Biophys. Acta 1829, 1147–1159.

Neuhold, L.A., and Wold, B. (1993). HLH forced dimers: tethering MyoD to E47 generates a dominant positive myogenic factor insulated from negative regulation by Id. Cell 74, 1033–1042.

Qiu, X., Mao, Q., Tang, Y., Wang, L., Chawla, R., Pliner, H., and Trapnell, C. (2017a). Reversed graph embedding resolves complex single-cell developmental trajectories.

Qiu, X., Hill, A., Packer, J., Lin, D., Ma, Y.-A., and Trapnell, C. (2017b). Single-cell mRNA quantification and differential analysis with Census. Nat. Methods 14, 309–315.

Rabiner, L., and Juang, B.H. (1993). Fundamentals of speech recognition (PTR Prentice Hall).

Salvatori, G., Lattanzi, L., Coletta, M., Aguanno, S., Vivarelli, E., Kelly, R., Ferrari, G., Harris, A.J., Mavilio, F., and Molinaro, M. (1995). Myogenic conversion of mammalian fibroblasts induced by differentiating muscle cells. J. Cell Sci. 108 (Pt 8), 2733–2739.

Sartori, R., Schirwis, E., Blaauw, B., Bortolanza, S., Zhao, J., Enzo, E., Stantzou, A., Mouisel, E., Toniolo, L., Ferry, A., et al. (2013). BMP signaling controls muscle mass. Nat. Genet. 45, 1309–1318.

Serra, C., Palacios, D., Mozzetta, C., Forcales, S.V., Morantte, I., Ripani, M., Jones, D.R., Du, K., Jhala, U.S., Simone, C., et al. (2007). Functional interdependence at the chromatin level between the MKK6/p38 and IGF1/PI3K/AKT pathways during muscle differentiation. Mol. Cell 28, 200–213.

Takahashi, K., and Yamanaka, S. (2006). Induction of pluripotent stem cells from mouse embryonic and adult fibroblast cultures by defined factors. Cell 126, 663–676.

Takahashi, K., Tanabe, K., Ohnuki, M., Narita, M., Sasaki, A., Yamamoto, M., Nakamura, M., Sutou, K., Osafune, K., and Yamanaka, S. (2014). Induction of pluripotency in human somatic cells via a transient state resembling primitive streak-like mesendoderm. Nat. Commun. 5, 3678.

Trapnell, C., Cacchiarelli, D., Grimsby, J., Pokharel, P., Li, S., Morse, M., Lennon, N.J., Livak, K.J., Mikkelsen, T.S., and Rinn, J.L. (2014). The dynamics and regulators of cell fate decisions are revealed by pseudotemporal ordering of single cells. Nat. Biotechnol. 32, 381–386.

Treutlein, B., Lee, Q.Y., Camp, J.G., Mall, M., Koh, W., Shariati, S.A.M., Sim, S., Neff, N.F., Skotheim, J.M., Wernig, M., et al. (2016). Dissecting direct reprogramming from fibroblast to neuron using single-cell RNA-seq. Nature 534, 391–395.

Vierbuchen, T., and Wernig, M. (2011). Direct lineage conversions: unnatural but useful? Nat. Biotechnol. 29, 892–907.

Vintsyuk, T.K. (1968). Speech discrimination by dynamic programming. Cybern. Syst. Anal. 4, 52–57.

Wang, D.Z., Valdez, M.R., McAnally, J., Richardson, J., and Olson, E.N. (2001). The Mef2c gene is a direct transcriptional target of myogenic bHLH and MEF2 proteins during skeletal muscle development. Development 128, 4623–4633.

Weintraub, H., Tapscott, S.J., Davis, R.L., Thayer, M.J., Adam, M.A., Lassar, A.B., and Miller, A.D. (1989). Activation of muscle-specific genes in pigment, nerve, fat, liver, and fibroblast cell lines by forced expression of MyoD. Proceedings of the National Academy of Sciences 86, 5434–5438.

Xu, J., Du, Y., and Deng, H. (2015). Direct Lineage Reprogramming: Strategies, Mechanisms, and Applications. Cell Stem Cell 16, 119–134.

Zaret, K.S., and Carroll, J.S. (2011). Pioneer transcription factors: establishing competence for gene expression. Genes Dev. 25, 2227–2241.

